# Cysteine reduction rather than methionine restriction promotes immunometabolic fitness to confer survival benefit in aging

**DOI:** 10.64898/2026.09.04.749519

**Authors:** Lucie Orliaguet, Hee-Hoon Kim, Piyush Gupta, Aileen Lee, Yuanjiu Lei, Ruiqi Li, Carisa Zeng, Yun-Hee Youm, Tamara Dlugos, Chenyang Guan, Andrew Wang, Maxim N. Artyomov, Yuval Kluger, Björn Schumacher, Vishwa Deep Dixit

## Abstract

Elevated sulfur-containing amino acids (SAA; methionine and cystine) are linked to human mortality. While methionine restriction (MR) extends lifespan in animals, it is often ignored that traditional longevity-promoting MR diets also lack cystine. When dietary cystine is eliminated, host redirects substrates from methionine cycle and generates cysteine via cystathionine γ-lyase (CTH) in transsulfuration pathway, leaving it unclear which SAA controls aging. Here, we show that selective cysteine restriction in *Caenorhabditis elegans* enhanced lifespan and stress survival independently of methionine. Notably, cysteine-free diets (methionine-replete or restricted) in *Cth-*deficient mice induced pro-metabolic effects, whereas MR with normal cysteine was ineffective. Sustained 70% cysteine reduction in aged *Cth^-^*^/-^ mice reprogrammed the immunometabolic axis, conferring healthspan benefits. Thus, lowering cysteine while keeping the methionine pool intact enhances healthy lifespan.

## INTRODUCTION

Cysteine is a dietary non-essential yet biochemically irreplaceable amino acid.^1^ Methionine, the other sole proteinogenic sulfur amino acid (SAA), lacks a thiol group and hence cannot form complexes with metals, including disulfide bond formation, nucleophilic catalysis, and redox signaling.^1,2^ Typically, cysteine and methionine contents are high in animal-based diets, while plant-based vegetarian foods are lower in SAA.^3^ Upon dietary protein intake, methionine is catabolized by methionine adenosyltransferase 2A (MAT2A) to generate S-adenosyl-methionine (SAM), a critical universal methyl donor.^1^ Methionine metabolism is a key node in organismal biology because methyltransferases catalyze methylation reactions that transfer methyl groups to both histone and nonhistone proteins. In the methionine cycle, glycine N-methyltransferase (GNMT) produces S-adenosyl-homocysteine (SAH) from SAM. Subsequently, the enzyme SAH hydrolase (also known as adenosylhomocysteinase) catalyzes the reversible hydrolysis of SAH to produce homocysteine. If the dietary cysteine is insufficient, the transsulfuration pathway (TSP) shuttles homocysteine to produce cystathionine, then cystathionine γ- lyase (CTH) hydrolyzes it into cysteine.^1^ The methionine salvage pathway, which replenishes methionine from SAM, produces spermidine, another established pro-longevity metabolite.^4^ Importantly, metabolomics data from humans who underwent sustained caloric restriction (CALERIE-II trial), which activates longevity mechanisms, demonstrated a decrease in the methionine cycle (reduction in Betaine-Homocysteine S-Methyltransferase (BHMT) and lower dimethylglycine), rewiring of TSP, reduction of cysteine and GSH, and elevation of taurine, a sulfonic non-synthetic amino acid.^5^ Thus, the metabolism of SAA, including rewiring of TSP and cysteine metabolism, is linked to improved metabolic health and lifespan extension in model organisms.

It is now recognized that cysteine is essential for organismal metabolism through pleiotropic effects, including regulation of CoA and GSH-mediated redox balance.^5,6^ Creation of forced cysteine depletion by deletion of CTH and feeding a cysteine-free (CysF) diet causes rapid, progressive 30% weight loss.^5,6^ In addition, cysteine depletion induces activation of central nervous system nuclei that control sympathetic nervous system (SNS)-derived norepinephrine, which causes rapid adipose tissue browning and thermogenesis through β3-adrenergic receptors.^5^ Notably, adipose browning has also been reported to be induced by methionine restriction.^7^ However, an important underappreciated issue is that methionine restriction is erroneously called MR because this diet also lacks cystine.^8,9^ Interestingly, it has been well established that an 80% reduction of methionine content (in a diet with 0% cystine) in adult rats resulted in robust 40% increases in median and maximum lifespan.^10,11^ Additional independent studies confirmed that MR, which in fact lacks both methionine and cysteine, robustly enhances lifespan in mice despite an increase in food intake, without restriction of calories.^12,13^ Moreover, using methionine-and cysteine-titrated diets in wild-type mice, the anti-adiposity effects of SAAR have been linked to cysteine restriction.^14^ However, when cysteine is restricted, the methionine cycle shuttles substrates into CTH-mediated cysteine synthesis in TSP, allowing the host to replenish cysteine and survive nutrient stress. Therefore, the specific effects of selective cysteine restriction cannot be delineated from methionine in wild-type animals. Moreover, given that cystine (oxidized dimeric form of cysteine) has been identified as the top upregulated metabolite linked to increased biological aging and mortality in humans^15^ and our recent findings that cysteine depletion induces pro-metabolic adipose thermogenesis,^5^ we tested the hypothesis that selective reduction of cysteine promotes metabolic health and lifespan.

## RESULTS

### Cysteine depletion, but not methionine restriction, drives mechanisms that reduce adiposity and enhance lifespan

We first investigated the effects of a traditional MR diet, which is actually a methionine and cysteine restriction (MCR) diet, on metabolic health across lifespan. Young and old wild-type male mice were fed an MCR (0.17% methionine, 0% cystine) or isocaloric control (CTRL: 0.86% methionine, 0.39% cystine) diet for 12 weeks (Figure 1A). Methionine is restricted to 0.17% because this level provides the minimum amount of this essential amino acid while maintaining methionine restriction. Consistent with previous findings identifying SAA restriction as a metabolically protective intervention throughout life,^10–13^ MCR prevented body weight gain in both young and old groups (Figure 1B). Compared to the CTRL diet, the MCR reduced the visceral adipose tissue (VAT) mass in old mice (Figure 1C), indicating a primary effect on adiposity. The MCR feeding improved systemic glucose homeostasis, with reduced fasting glycemia in old mice and improved glucose and insulin tolerance in young and old mice (Figures 1D, 1E, and S1A-S1C). In addition, MCR also protected from age-related increases in hepatic steatosis (Figure 1F), thus confirming prior studies that MCR improves metabolic health. We next examined whether the absence of cystine in the MCR diet impacts the TSP enzyme CTH, which is required for cysteine generation. Notably, absence of cystine in the MCR diet caused an upregulation of hepatic CTH in both young and old mice (Figure 1G), suggesting activation of TSP to synthesize cysteine.

**Figure 1.**
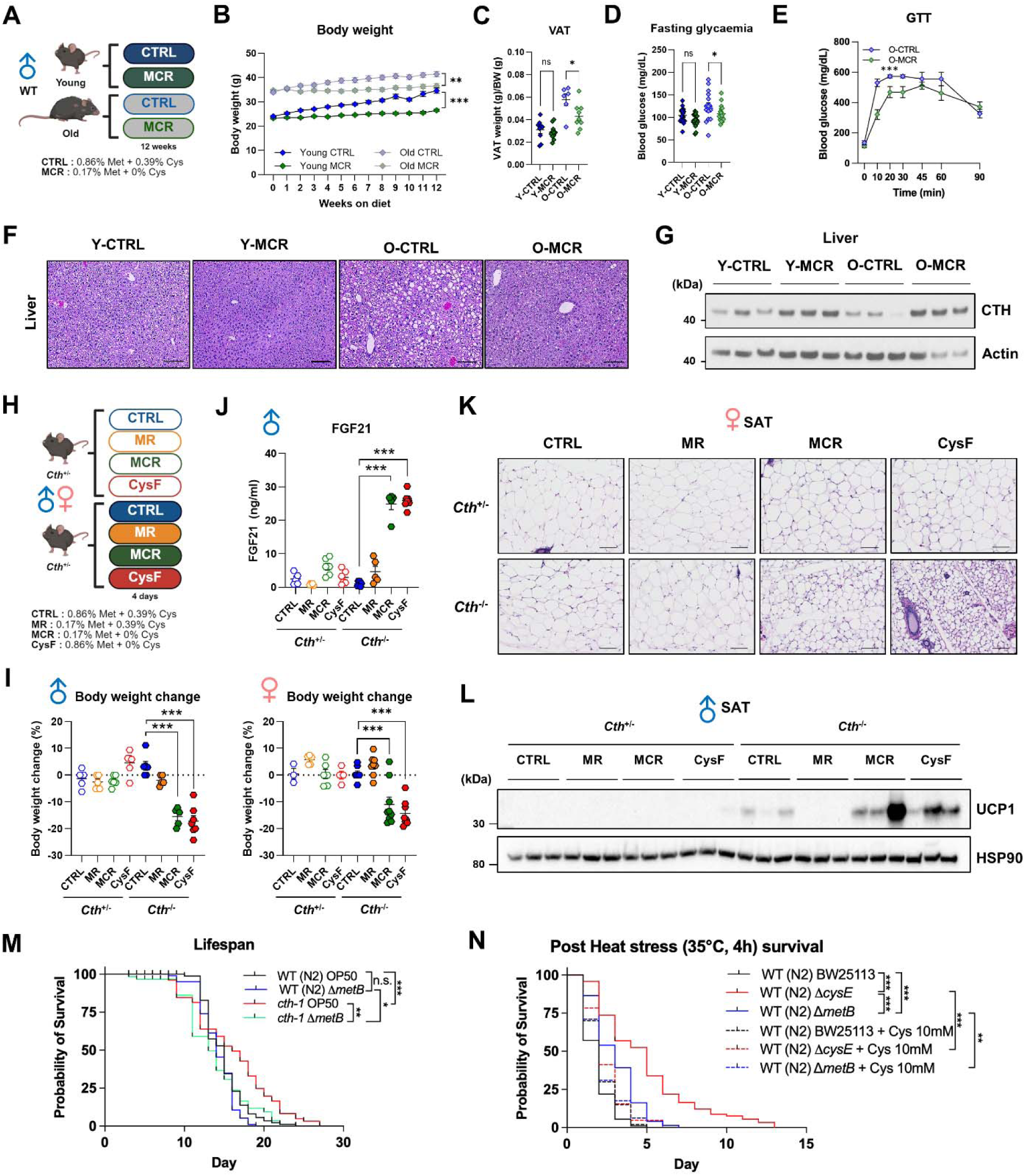
Cysteine depletion, but not methionine restriction, drives mechanisms that reduce adiposity and enhance lifespan. (A-G) Young or old male mice were fed either the control diet (CTRL) or the methionine-and cysteine-restricted diet (MCR) for 12 weeks (*n* = 15-21/group). (A) Schematic outline with diet compositions. (B) Body weight changes. (C) Visceral adipose tissue (VAT) weight index (*n* = 6-11/group). (D) Fasting blood glucose levels (*n* = 17-21/group). (E) Glucose tolerance test in old mice (*n* = 4/group). (F) Representative H&E staining of liver tissues. (G) Representative CTH western blot (3 blots/group) of liver tissues. (H-L) Adult *Cth*^+/-^ and *Cth*^-/-^ mice were fed a CTRL, methionine-restricted (MR), MCR, or cysteine-free (CysF) diet for 4 days (*n* = 5-7/group for male and 3-9 for female mice). (H) Schematic outline with diet compositions. (I) Body weight change over 4 days in male (left) and female (right) mice. (J) Serum FGF21 levels in male mice. (K) Representative H&E staining of subcutaneous adipose tissue (SAT) of female mice. Scale bars = 50 µm. (L) Representative UCP1 western blot (3 blots/group) of SAT of male mice. (M) Lifespan assays were performed in WT (N2) and *cth*-*1(ok3319)V C. elegans* fed an OP50 (wild-type) or methionine-depleted (Δ*metB*) *E. coli* strain (*n* = 74-107/group). (N) WT (N2) *C. elegans* were fed a wild-type (BW25113), cysteine-depleted (Δ*cysE*), or methionine-depleted (Δ*metB*) *E. coli* strain with or without 10 mM cysteine from the L1 stage to day 1 adulthood, followed by heat stress (35 °C) exposure for 4 h. Survival was analyzed (*n* = 74-100/group). Data are presented as mean±SEM. Unpaired two-tailed t-tests (B-E), Two-way ANOVA with Sidak’s correction for multiple comparisons (I and J), or two-sided log-rank (Mantel-Cox) hypothesis test (M and N) were performed. Figures 1A and 1H were created with Biorender.com. *P <0.05, **P <0.01, and ***P <0.001. ns, not significant.

We next sought to disentangle the respective contributions of methionine and cysteine in SAA restriction-induced metabolic effects. To do this, we used *Cth^+/-^*and CTH–deficient (*Cth*⁻^/^⁻) mice fed isocaloric diets selectively restricting methionine and/or cysteine (Control diet (CTRL: 0.86% methionine, 0,39% cystine); methionine restricted (MR: 0.17% methionine, with normal 0.39% cystine); methionine and cysteine restricted (MCR: 0.17% methionine, with 0% cystine); or cystine-free (CysF: normal 0.86% methionine, 0% cystine)) (Figure 1H). Strikingly, in *Cth^+/-^* mice, which are similar to wild-type mice, the MR did not significantly reduce body weight, but only cystine-restricted conditions (*Cth*^-/-^ mice fed CysF or MCR diets) induced rapid body weight loss in both sexes (Figure 1I). Consistent with our previous findings,^5^ this body weight loss was attributable to reduced fat mass and occurred without changes in food intake (Figures S1D-S1F).

The central driver of the beneficial effects of MCR is fibroblast growth factor 21 (FGF21), a pro-longevity hepatokine.^7,16,17^ Through generation of FGF21 and CTH double knockout mice, we recently demonstrated that despite robust induction of FGF21 by cysteine depletion in mice, there was only partial protection from weight loss, and deletion of FGF21 did not affect adipose tissue browning.^5^ Therefore, we investigated the impact of specific SAA-deficient diets on FGF21 production. Interestingly, in both *Cth*^+/-^ and *Cth*^-/-^ mice, MR did not induce FGF21, whereas the MCR and CysF diets strongly induced FGF21 in both sexes of *Cth*^-/-^ mice, where TSP is made essential (Figures 1J and S1G). These data demonstrate that cysteine but not methionine loss drives neuroendocrine programming that drives FGF21 production. Growth differentiation factor 15 (GDF15) is another metabolic hormone induced in response to nutritional stress.^18^ However, its contribution to the improvement of metabolic health in the context of MCR is unknown. Similar to FGF21 and a previous 4-day cysteine restriction model,^5^ we found increased GDF15 levels in *Cth*^-/-^ mice fed with MCR and CysF compared to those of the *Cth*^+/-^ mice, and it was not sufficient to significantly impact food intake (Figures S1F and S1H).

Beige adipocytes serve as a ‘catabolic sink’ for glucose, lipids, and branched-chain amino acids, clearing these metabolites from the circulation with uncoupled thermogenesis to convert chemical energy into heat.^19,20^ Comparative transcriptomic analyses indicate that inducible thermogenic fat in human adults is similar to murine beige fat,^21^ making it an important target in aging. Given that both MR and cysteine depletion induce adipose tissue thermogenesis and uncoupling, we next tested the contribution of methionine or cysteine to adipose tissue beiging. Histological analyses of the three different adipose depots—brown adipose tissue (BAT), subcutaneous adipose tissue (SAT), and VAT— revealed a profound remodeling of adipose structure when cysteine but not methionine was restricted, with a substantial beiging in SAT, characterized by smaller adipocytes with multilocular lipid droplets, and reduced lipid accumulation in BAT (Figures 1K and S1I). VAT structure was unchanged across regimens (Figure S1I). In addition, the MCR and CysF diets fed to *Cth*^-/-^ mice also increased uncoupling protein 1 (UCP1) in SAT, whereas MR did not affect adipose tissue browning or UCP1 protein levels in control or *Cth*^-/-^ mice (Figure 1L and Figure S1I).

The TSP is evolutionarily conserved from *Caenorhabditis elegans* to humans.^1,2^ Therefore, we next utilized *C. elegans* lacking the ability to synthesize cysteine (*cth-1*) to determine cysteine’s effects on lifespan and stress resilience. Interestingly, compared to control wild-type (WT (N2)) worms fed with normal OP50 *Escherichia coli*, the *cth-1* mutants lived significantly longer (Figure 1M). However, dietary methionine restriction through feeding with a *metB* mutant *E. coli*, which does not synthesize methionine, did not extend lifespan in control WT (N2) and rather significantly reduced lifespan in *cth-1* mutant *C*. *elegans* (Figure 1M). We further tested the necessity of cysteine in host physiological resilience by utilizing a heat stress-induced mortality model. The ability to survive heat stress is a typical longevity-associated resilience phenotype in nematodes.^22^ As expected, heat stress induced rapid mortality in WT (N2) *C. elegans* seeded with normal BW25113 *E. coli* (Figure 1N). Notably, both dietary methionine and cysteine restriction through engineered *E. coli* that cannot synthesize methionine (Δ*metB*) or cysteine (Δ*cysE*) significantly extended post-heat stress survival in *C. elegans*, where cysteine restriction (Δ*cysE*) showed a significantly stronger effect than methionine restriction (Δ*metB*) (Figure 1N). Moreover, the lifespan extension effect of cysteine restriction was abrogated when cysteine was restored in cysteine-restricted *C. elegans* (Figure 1N), suggesting that cysteine but not methionine restriction drives organismal fitness and healthy aging.

### Cysteine depletion retains adipose beiging potential in aged mice

It is presently unclear whether adipose tissue in advanced age is amenable to metabolic reprogramming that induces thermogenesis.^20^ Given that *C. elegans* lacks the adipose tissue thermogenic machinery, we next tested whether mechanisms that promote adipose tissue beiging are regulated by cysteine in aged mice. To test this, we aged the control and *Cth*^⁻/⁻^ mice up to 23 months and challenged them with a CysF diet for 5 days (Figure 2A). Interestingly, cysteine depletion induced rapid, significant 20% body weight loss within 5 days in aged *Cth*^⁻/⁻^ mice, although the magnitude of body weight loss was lower than in young mice (Figure 2B). Notably, cysteine depletion in aged mice induced rapid adipose browning and reduced lipid accumulation in the liver without affecting food intake and grip strength (Figures 2C-2E). As previously reported in adult mice,^5^ cysteine depletion in old animals caused a robust increase in circulating FGF21 concentrations (Figure 2F). Moreover, consistent with reduced lipid deposition, compared to control age-matched littermates, the BAT showed gene expression profiles that suggest increased thermogenesis and lipid turnover (Figure 2G). Furthermore, cysteine depletion affected liver metabolic regulators, with a decrease in genes associated with lipid synthesis (*Fasn*) and an increase in a lipolysis-related gene (*Pnpla2*) (Figure 2H). Interestingly, *Igfbp2*, which encodes insulin-like growth factor-binding protein 2 (IGFBP2), an endogenous protector against liver steatosis,^23^ was significantly increased by cysteine depletion (Figure 2H). Consistent with increased thermogenesis, the SAT of cysteine-depleted aged mice had an increase in *Cox7a1*, *Cidea*, *Elovl3,* and *Slc27a2* mRNA as well as elevation of UCP1 protein (Figures 2I and 2J). These data suggest that cysteine is the primary driver of metabolic reprogramming in aging.

**Figure 2.**
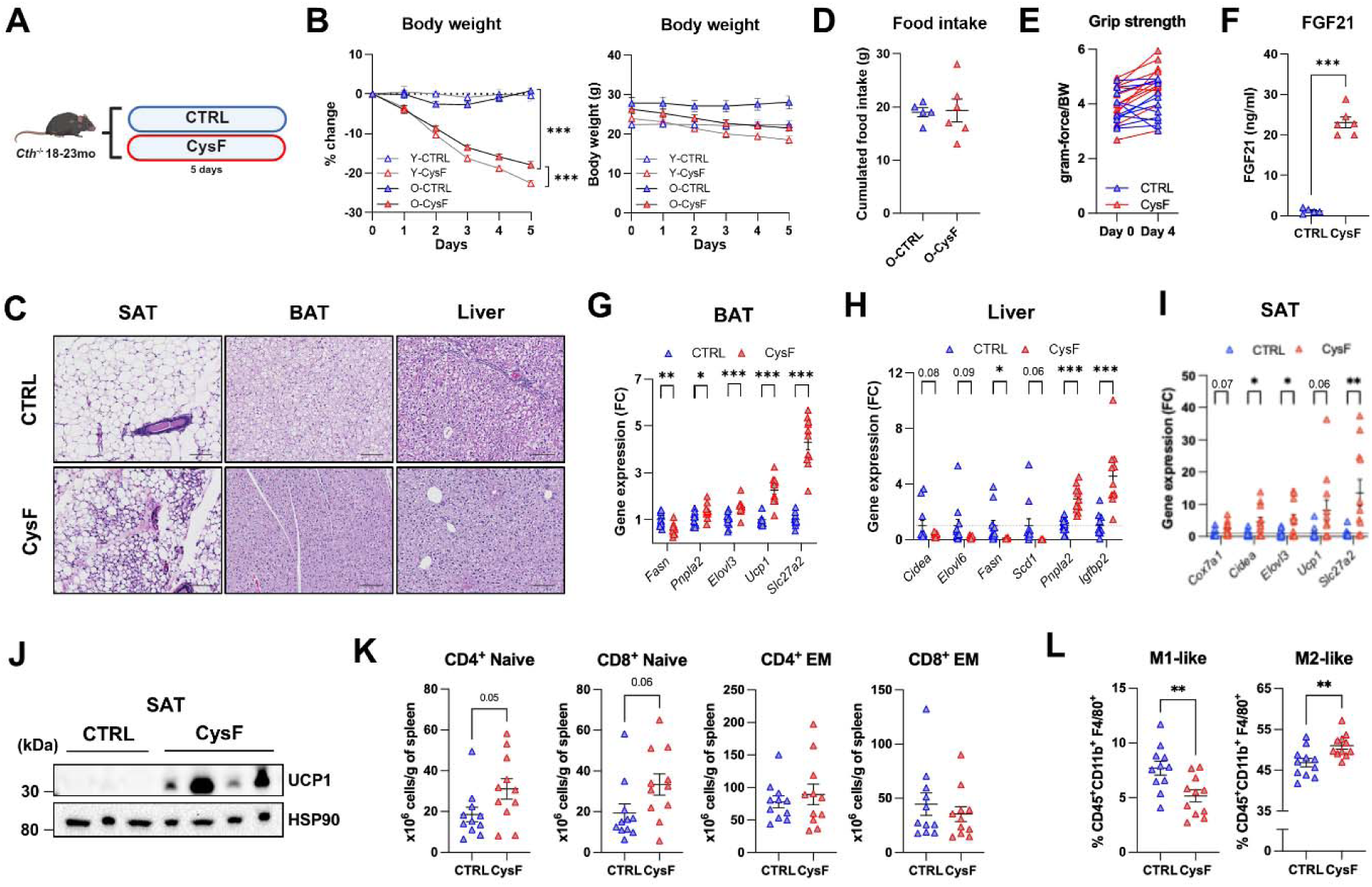
Cysteine depletion retains adipose beiging potential in aged mice. (A-L) 18-to 23-month-old *Cth*^-/-^ mice were fed a CTRL or CysF diet for 5 days (*n* = 11/group). (A) Schematic outline. (B) Body weight changes per proportion (left) or per gram (right) of both young (*n* = 10/group) and old *Cth*^-/-^ mice fed a CTRL or CysF diet. (C) Representative H&E staining of SAT, BAT, and liver tissues. Scale bars = 50 µm. (D) Cumulative food intake for 5 days (*n* = 5-6/group). (E) Longitudinal grip strength measurement. (F) Serum FGF21 levels (*n* = 5-6/group). (G-I) qRT-PCR analyses for BAT (G), Liver (H), and SAT (I). (J) Representative UCP1 western blot (3 blots/group) of SAT. (K and L) Flow cytometry analyses of CD4^+^ and CD8^+^ naïve and effector/memory (EM) T cells (K) and M1-and M2-like macrophages (L). Data are presented as mean±SEM. Unpaired two-tailed t-tests (D-L) or Two-way ANOVA with Sidak’s correction for multiple comparisons (B) were performed. Figure 2A was created with Biorender.com. *P <0.05, **P <0.01, and ***P <0.001.

As aging profoundly impairs immune function,^24^ we tested whether cysteine-induced immunometabolic reprogramming can lower inflammaging. One of the hallmark features of aging of the adaptive immune system in mice and humans is loss of naïve T cells and sustained activation of the innate immune system.^24^ Interestingly, cysteine depletion in old mice tended to increase both splenic CD4^+^ and CD8^+^ naïve T cells, without altering effector-memory T cell subsets or total splenocyte counts (Figures 2K, S1J, and S1K). Consistent with this, we found that cysteine depletion significantly decreased the frequencies of pro-inflammatory M1-like (CD11c^+^) macrophages and increased reparative M2-like (CD206^+^) macrophages in the spleen of old mice (Figures 2L and S1L). Together, these data demonstrate that short-term cysteine depletion induces rapid beneficial metabolic remodeling in old mice, improving adipose and hepatic function, and lowering several hallmarks of immunological aging.

### Sustained cysteine reduction is sufficient to promote metabolic health during aging

Complete cysteine depletion in mice (*Cth^-/-^* on a CysF diet) triggers unchecked thermogenesis at subthermoneutral and thermoneutral temperatures, leading to rapid 30% weight loss, requiring euthanasia.^5^ This indicates that cysteine is essential for host survival.^5,6^ Therefore, we tested whether moderate and sustained cysteine restriction (CysR) can be harnessed and if it retains metabolic benefits as a potential intervention. We previously reported that in cysteine-depleted mice, restoration of 75% cysteine content of a normal diet fully rescued weight loss.^5^ Given that a reduction of 25% dietary cystine content in *Cth^-/-^* mice had no impact on weight loss (Figure 3A), we further increased the reduction of cystine concentration to 70%. The 70% CysR (0.12% cystine) in young adult *Cth^-/-^* mice induced 10% body weight loss within the first two weeks of the intervention, after which body weight stabilized for the duration of the 11-week treatment (Figures 3B, 3C, and S2A). Of note, the 70% CysR group consumed more food than controls, possibly a compensatory response to reduced cystine intake (Figure 3D). We found that 70% CysR maintained glucose tolerance and performance on the running wheel and rotarod in adult mice (Figures S2B-S2D). In addition, during the weight-stabilized phase at the end of intervention, the *Cth^-/-^* mice on CTRL or 70% CysR showed no differences in total energy expenditure (Figure S2E). Notably, 70% CysR induced FGF21 production (Figure 3E) to a similar magnitude as seen in the CysF diet in *Cth^-/-^* mice (Figure 1J), which showed rapid 30% weight loss. Thus, it appears that induction of FGF21 serves as a biomarker of cysteine reduction-induced metabolic stress response rather than a driver of weight loss.

**Figure 3.**
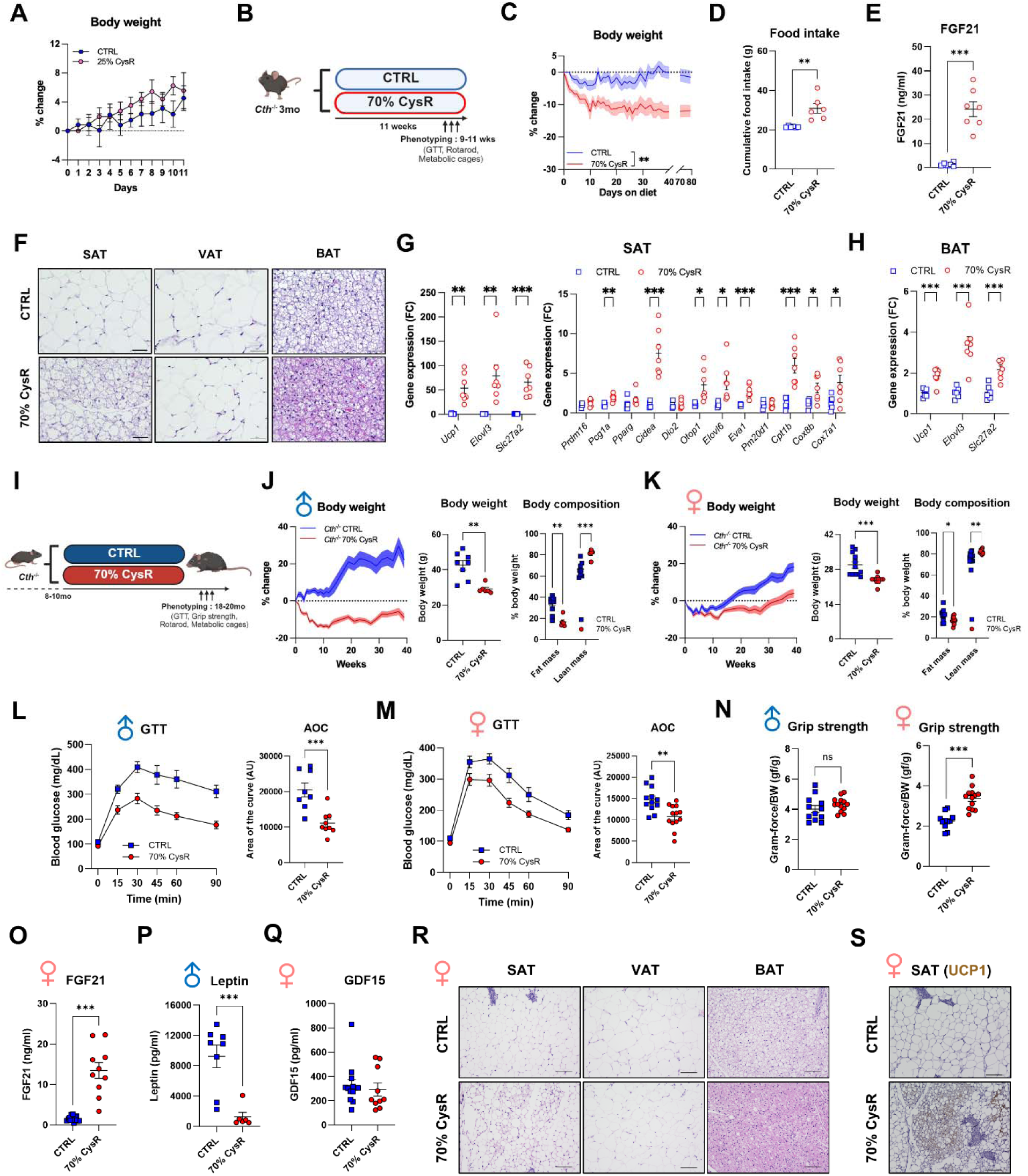
Sustained cysteine restriction is sufficient to promote metabolic health during aging. (A) 3-month-old *Cth*^-/-^ male mice were fed a CTRL or 25% cysteine-restricted diet for 11 days (*n* = 6/group). (B-H) 3-month-old *Cth*^-/-^ mice were fed a CTRL or 70% CysR diet for 11 weeks (*n* = 6-7/group). (B) Schematic outline. (C) Body weight. (D) Cumulative food intake for 5 days (*n* = 6/group. (E) Serum FGF21 levels. (F) Representative H&E staining of SAT, VAT, and BAT. (G and H) qRT-PCR analyses for SAT (G) and BAT (H). (I-S) 8-to 10-month-old *Cth*^-/-^ mice were fed a CTRL or 70% CysR diet for 10 months (*n* = 13-25/group). (I) Schematic outline. (J and K) Body weight changes per proportion (left), per gram (middle), and body composition analysis (right) in male (J) and female (K) mice. (L and M) GTT in male (L) and female (M) mice (*n* = 8-13/group). (N) Grip strength tests of male (left) and female (right) mice (*n* = 11-13/group). (O) Serum FGF21 levels in female mice (*n* = 10-13/group). (P) Serum leptin levels in male mice (*n* = 6-8/group). (Q) Serum GDF15 levels in female mice (*n* = 10-13/group). (R) Representative H&E staining of SAT, VAT, and BAT of female mice. (S) Representative UCP1 immunohistochemistry in SAT of female mice. Scale bars = 50 µm. Data are presented as mean±SEM. Unpaired two-tailed t-tests were performed. Figures 3B and 3I were created with Biorender.com. *P <0.05, **P <0.01, ***P <0.001, and ****P <0.0001. ns, not significant.

Interestingly, at the tissue level, moderate 70% CysR restriction induced selective adipose remodeling. The SAT of *Cth^-/-^* mice on 70% CysR exhibited moderate patches of browning, while VAT showed reduced adipocyte size without browning (Figure 3F). In addition, BAT of 70% CysR mice displayed enhanced thermogenic features, with less lipid accumulation (Figure 3F). Consistent with the adipose browning response, the SAT and BAT showed increased expression of canonical thermogenic and browning markers, including *Ucp1*, *Elovl3*, and *Slc27a2* (Figures 3G and 3H). Similarly, after moderate CysR, *Ppargc1a*, *Cidea*, *Otop1*, *Elovl6*, *Cpt1a*, *Cox8b* and *Cox7a1* expression was significantly induced in SAT, and *Otop1* and *Eva1* in BAT (Figures 3G and S2F). In contrast, these thermogenic genes were not induced in VAT (Figure S2G). Together, these data show that moderate CysR is sustainable and induces beneficial and selective remodeling of SAT and BAT, activating thermogenic programs without compromising muscular function and activity during aging.

### Chronic cysteine restriction protects against metabolic dysfunction during aging

We next asked whether chronic CysR may improve metabolic health during aging without inducing potential trade-offs. To address this, CysR was started in *Cth^-/-^* mice from midlife (8-10 months old) to old age (up to 20 months old), when comprehensive phenotyping was performed (Figure 3I). Importantly, 70% CysR in *Cth^-/-^* mice, which cannot perform TSP to generate endogenous cysteine, protected against age-related weight gain and loss of lean mass and improved glucose tolerance in both sexes, whereas grip strength was improved only in male mice and food intake and performance on rotarod were comparable (Figures 3J-3N and S2H-S2J). In addition, long-term 70% CysR robustly enhanced FGF21 and reduced leptin (Figures 3O and 3P), which is consistent with reduced fat mass. Notably, unlike complete cysteine depletion that elevates GDF15, 70% CysR did not impact the circulating concentration of GDF15 (Figure 3Q). Moreover, 70% chronic CysR in aged mice induced browning of SAT, reduced adipocyte size in VAT, and lowered lipid accumulation in BAT (Figure 3R). In particular, SAT of 70% CysR mice exhibited increased UCP1 expression and reduced adipocyte size (Figures 3R, 3S, and S2K), which may all contribute to better insulin sensitivity in old *Cth*^-/-^ mice upon 70% CysR feeding. Using indirect calorimetry, we investigated whether changes in energy expenditure and respiratory exchange ratios (RER) contribute to the beneficial metabolic effects. With the data represented as a regression between energy expenditure and body weight, 70% CysR mice exhibit a significantly steeper slope and comparable absolute energy expenditure despite lower body weight (Figure S2L). RER did not differ between groups when the body weight had stabilized (Figure S2M), suggesting that this altered energy expenditure–body mass relationship was not accompanied by a detectable shift in substrate utilization. Together with the increased expression of thermogenic markers, such as UCP1, and enhanced adipose tissue beiging, these findings are consistent with increased thermogenic capacity and metabolic responsiveness in 70% CysR mice.

Of note, we applied the same long-term 70% CysR regimen to assess the effects of a moderate CysR in littermate wild-type (*Cth*^+/+^) mice, which possess an intact TSP (Figure S3A). In wild-type mice, 70% CysR didn’t affect body weight gain, muscle function, and glucose homeostasis (Figures S3B-S3F). Of note, fat mass was increased in male mice upon cysteine restriction (Figure S3B). In addition, 70% CysR in wild-type mice did not induce browning of adipose depots, reduce liver lipid content, or impact serum tumor necrosis factor (TNF)-α, monocyte chemoattractant protein-1 (MCP-1), interleukin (IL)-6, and leptin levels (Figures S3G and S3H). Together, these findings indicate that long-term moderate CysR in *Cth^-/-^*mice, but not in wild-type mice, improves metabolic health during aging without adverse physiological trade-offs.

### Chronic cysteine restriction induces metabolic reprogramming that favors healthspan

As long-term moderate CysR improves adipose tissue beiging and metabolic functions, we investigated the induction of transcriptional programs in SAT and BAT using single-nucleus RNA sequencing (snRNA-seq) of SAT and BAT depots from *Cth^-/-^* mice fed a CTRL or 70% CysR diet. A total of 13,353 (CTRL) and 12,901 (70% CysR) nuclei from SAT and 42,135 (CTRL) and 34,786 (70% CysR) nuclei from BAT were analyzed (Figures S4A and S4B). Unbiased clustering identified adipose cell types,^25^ namely adipocytes, adipose stem cells (ASC), immune cells, and endothelial cells in SAT and BAT, epithelial cells and smooth muscle cells in SAT only, and myocytes in BAT only (Figures S4A-S4D). In both depots, cysteine restriction reprogrammed the cell state and transcriptomic response, rather than cellular proportions, except for a higher mature adipocyte fraction in BAT of 70% CysR mice compared to controls (Figures S4A and S4B). Therefore, we focused our downstream analysis on the adipocyte compartment.

Unbiased clustering of SAT adipocyte fraction identified 5 distinct clusters, each defined by a specific transcriptional program: SA0, basal white adipocyte markers (*Fabp4*, *Lipe*, *Adipoq*, *Plin4* and *Nnat*); SA1, extracellular matrix remodeling (*Serpine1*, *Col5a1*, *Col6a3*); SA2, lipogenesis (*Acly*, *Acaca*, *Elovl6* and *Fgf14*); SA3, beige/brown associated genes (*Ucp1* and *Cidea*); and SA4, interferon response (*Ifit3*, *Ifi44* and *Oasl2*) (Figures 4A and 4B). Interestingly, consistent with histological analysis, we found that 70% CysR induced expansion of the beige (SA3) adipocytes (Figure 4A). Accordingly, gene set enrichment analysis (GSEA) showed that 70% CysR upregulated thermogenesis-and oxidative phosphorylation (OXPHOS)-related genes in overall adipocytes (Figures 4C and S4E). Moreover, SA3 beige adipocytes in 70% CysR mice exhibited significantly increased expression levels of genes related to fatty acid metabolic processes, the electron transport chain, and fatty acid and lipid oxidation (Figure 4D), indicating that moderate CysR enhanced not only beige cell abundance but also the thermogenic potential of these cells.

**Figure 4.**
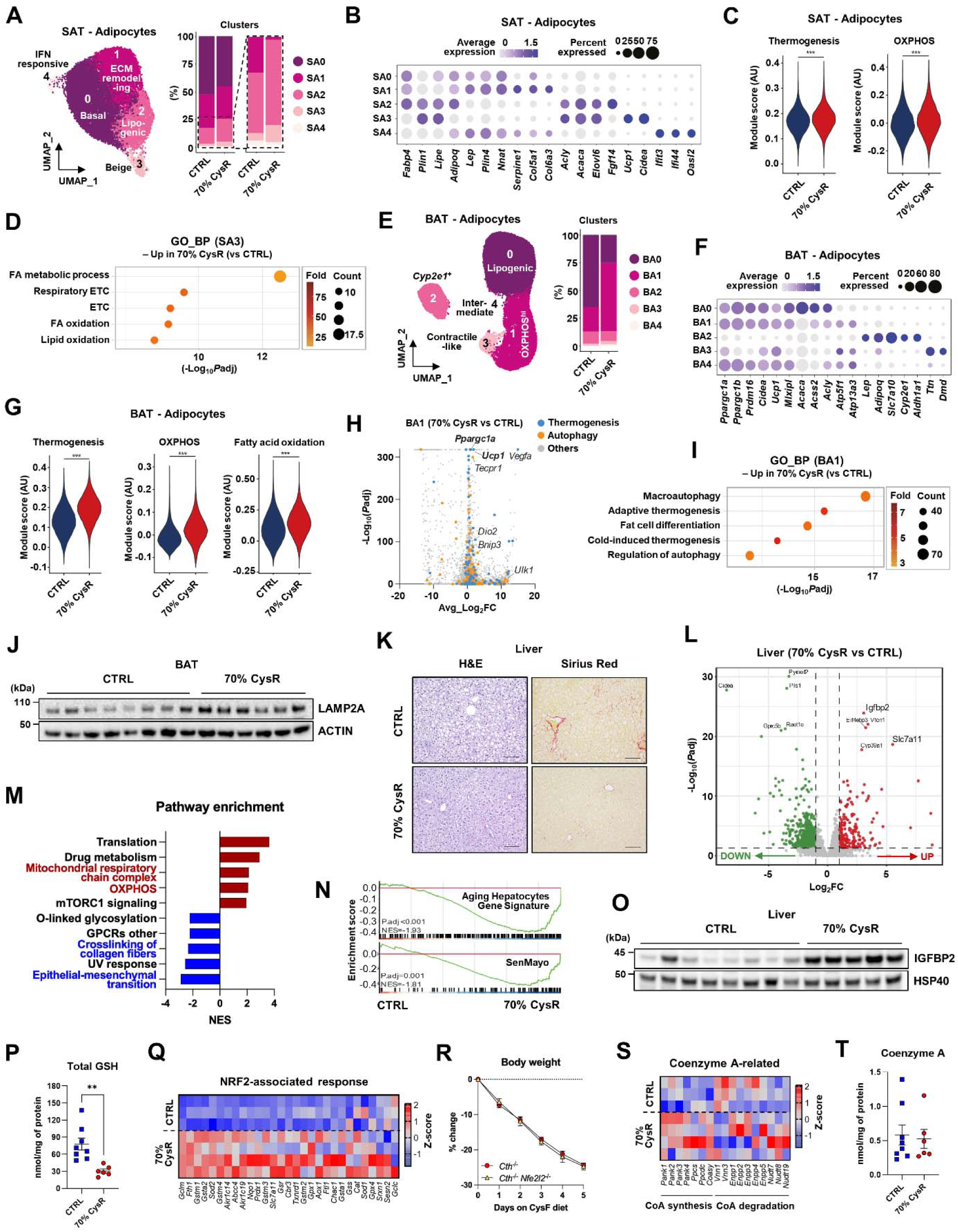
Chronic cysteine restriction induces metabolic reprogramming that favors healthspan. (A-D) Single-nucleus RNA sequencing (snRNA-seq) of SAT from 8-to 10-month-old *Cth*^-/-^ mice fed a CTRL or 70% CysR diet for 10 months. (A) UMAP for the SAT adipocyte subclusters. (B) Marker genes for each cluster. (C) Scoring analysis for thermogenesis and OXPHOS pathways in the whole SAT adipocyte fraction. (D) Gene ontology (GO) biological pathway (BP) analysis for SA3 beige adipocytes. (E-I) SnRNA-seq of BAT from 8-to 10-month-old *Cth*^-/-^ mice fed a CTRL or 70% CysR diet for 10 months. (E) UMAP for the BAT adipocyte subclusters. (F) Marker genes for each cluster. (G) Scoring analysis for thermogenesis, OXPHOS, and fatty acid oxidation pathways in the whole BAT adipocyte fraction. (H) Volcano plot for differentially expressed genes in BA1 OXPHOS^hi^ adipocytes. (I) GO_BP analysis for BA1 OXPHOS^hi^ adipocytes. (J) Representative LAMP2A western blot of BAT (8 or 6 blots/group). (K) Representative H&E and Sirius Red staining of liver tissues. Scale bars = 50 µm. (L) Volcano plot for differentially expressed genes in the liver, determined by bulk RNA-seq (*n* = 3-4/group). (M) Pathway enrichment analysis. (N) Gene set enrichment analysis for indicated pathways in the liver. (O) Representative IGFBP2 western blot of the liver (8 or 5 blots/group). (P) Total GSH levels in the liver (*n* = 6-8/group). (Q) Heatmap for the genes involved in the NRF2-associated response from the liver RNA-seq. (R) *Cth*^-/-^ and *Cth*^-/-^*Nfe2l2*^-/-^ mice (*n* = 7-13/group) were fed a CysF diet for 5 days. Body weight was recorded every day. (S) Heatmap of the genes involved in the coenzyme A-related pathway from the liver RNA-seq. (T) Coenzyme A levels in the liver (*n* = 6-8/group). Data are presented as mean±SEM. Unpaired two-tailed t-tests were performed. **P <0.01 and ***P <0.001.

Then, we adopted the same strategy to assess BAT adipocytes. Similar to SAT, unbiased clustering of BAT adipocytes identified 5 different clusters: BA0, lipogenesis (*Acaca*, *Acly* and *Acss2*); BA1, high OXPHOS; BA2, *Cyp2e1*^+^ that suppresses thermogenesis^26^; BA3, contractile-like (*Ttn* and *Dmd*); and BA4, intermediate that shares characteristics of BA0 and BA1 (Figures 4E and 4F). Strikingly, moderate CysR induced a shift in adipocyte composition with the decrease of lipogenic BA0 and expansion of OXPHOS^hi^ BA1 cluster (Figure 4E). The reduction in lipogenic adipocytes (BA0) may contribute to attenuation of BAT involution, consistent with the preservation of a more thermogenic BAT phenotype.^27,28^ GSEA showed that BAT adipocytes in 70% CysR mice had significantly higher expression levels of genes related to thermogenesis, OXPHOS, and fatty acid oxidation across clusters compared to controls (Figures 4G and S4F). We next analyzed differentially expressed genes and pathways of the OXPHOS^hi^ BA1 cluster, which was most remarkably changed in proportion by 70% CysR. Interestingly, 70% CysR significantly induced not only thermogenesis-related genes, including *Ucp1*, but also various autophagy-related genes in the BA1 cluster (Figures 4H and 4I), suggesting that in addition to the proportional change, these cells may possess a higher per-cell basis thermogenic capacity than controls. Moreover, the upregulation of autophagy-related genes by 70% CysR was observed across adipocyte clusters (Figures S4G and S4H). Considering that activation of chaperone-mediated autophagy (CMA) restores aging-associated functional declines of BAT,^29^ we examined the expression levels of LAMP2A, the rate-limiting CMA component. We found that LAMP2A protein expression in BAT was significantly increased by 70% CysR (Figures 4J and S4I). Together, the coordinated response upon 70% CysR, in both SAT and BAT, may suggest that moderate CysR favors adipocyte profiles associated with metabolic flexibility and healthy aging.

### Cysteine restriction remodels the hepatic transcriptome and prevents age-associated liver steatosis

The liver is known for the highest expression of the CTH enzyme; however, liver-specific CTH deletion is not sufficient to reduce systemic cysteine levels and induce weight loss.^5^ As short-term and systemic cysteine depletion in old *Cth^-/-^* mice protected against age-associated liver steatosis (Figure 2C), we tested whether moderate CysR prevents age-related liver steatosis. Histological analysis of liver sections revealed a marked reduction in lipid droplet accumulation as well as decreased age-related fibrosis by 70% CysR (Figure 4K). To identify the transcriptional basis of this improvement, we performed bulk RNA-seq on the liver tissues. We found that CysR extensively remodeled the hepatic transcriptome, including mitochondrial function and OXPHOS pathways, and downregulated collagen fiber crosslinking and epithelial-mesenchymal transition pathways (Figures 4L, 4M, and S4J), the latter of which is a hallmark of aging.^30^ In addition, consistent with reduced hepatic steatosis, 70% CysR suppressed expression of genes related to lipid biosynthetic processes (Figure S4K). Moreover, age-associated hepatocyte gene signature^31^ and cellular senescence-like genes (SenMayo^32^) were significantly downregulated by 70% CysR (Figure 4N). A detailed differential gene expression analysis identified that *Igfbp2* was the most substantially induced gene by 70% CysR, which was also significantly upregulated at the protein level (Figures 4L, 4O, and S4L). Given that IGFBP2 is a negative regulator of IGF signaling and its expression level is inversely associated with obesity, type-2 diabetes, and hepatic steatosis,^23,33^ the metabolic benefits of 70% CysR may be in part mediated by upregulation of IGFBP2. In addition to *Igfbp2*, the gene expression level of *Slc7a11*, which encodes cystine-glutamate antiporter xCT, was significantly upregulated in 70% CysR mice (Figure 4L), indicating a compensatory response to uptake more cystine to maintain the cellular cystine pool.

Cysteine is a crucial component of the cellular antioxidant glutathione (GSH).^1^ Since we have previously shown that complete cysteine depletion reduces tissue glutathione levels,^5^ we assessed GSH-related oxidative stress pathways. While hepatic GSH levels were significantly decreased in 70% CysR mice, lipid peroxidation, assessed by 4-hydroxynonenal (4-HNE) protein levels, was comparable between the two groups (Figures 4P and S4M). Consistently, we observed an induction of nuclear factor erythroid 2-related factor 2 (NRF2) target genes, including *Gclm* and *Nqo1* (Figure 4Q), potentially as a compensatory antioxidant mechanism. These results suggest that redox balance was preserved in the 70% CysR condition. To study the possibility that NRF2 is required for CysF-induced body weight loss, we generated CTH and NRF2 double knockout mice (*Cth*^-/-^*Nfe2l2*^-/-^) and fed them a CysF diet. We found that *Cth*^-/-^*Nfe2l2*^-/-^ mice, upon complete cysteine depletion, lose weight similarly to *Cth*^-/-^ mice (Figure 4R), indicating that NRF2 is dispensable for cysteine depletion-induced weight loss and associated metabolic improvements.

Cysteine starvation has been reported to reduce coenzyme A (CoA) during weight loss.^5,6^ Despite the moderate weight loss and protection from subsequent weight gain through lifespan in 70% CysR mice, there were no significant differences in genes associated with CoA metabolism and hepatic CoA concentrations (Figures 4S and 4T), suggesting that CoA is unlikely to drive metabolic benefits in aging mice upon cysteine restriction. Additionally, cysteine contributes to the synthesis and maintenance of iron-sulfur clusters.^34^ Surprisingly, there was a trend of increase in the expression of genes associated with iron-sulfur cluster synthesis upon 70% CysR, suggesting an adaptive response to leverage the cysteine pool for maintenance of iron-sulfur cluster homeostasis (Figure S4N). Furthermore, cellular or lysosomal cysteine and cystine scarcity triggers the integrated stress response (ISR) and ferroptosis, respectively, by upregulating activating transcription factor 4 (ATF4).^35,36^ We found that the gene expression levels of *Ddit3*, a downstream target of ATF4 in ISR, and ferroptosis drivers and suppressors^37^ were unchanged in the liver of 70% CysR mice (Figures S4O and S4P). Together, these data suggest that sustained 70% CysR in aged mice induces mechanisms associated with fatty acid oxidation, thermogenesis, autophagy, and reduced IGF signaling, which have been linked to enhancing healthspan and lifespan.^38–40^

## DISCUSSION

Over the last few decades, SAA restriction has been extensively studied as a powerful pro-longevity intervention that extends healthspan and lifespan.^8–14^ In the present study, we identify cysteine, rather than methionine, as the nutritional determinant of the canonical benefits of SAA restriction on metabolic healthspan and lifespan in *C. elegans*. In particular, we utilized a combination of a transgenic mouse model (*Cth*^-/-^) with a selective CysF or CysR diet to experimentally uncouple methionine from cysteine. From these well-controlled experiments, we demonstrate that MR alone was ineffective, whereas CysR fully recapitulated the canonical SAA restriction phenotype, characterized by reduced adiposity and increased adipose browning. Our findings are consistent with a previous study showing that cysteine supplementation reverses the beneficial effects of SAA restriction on body weight and adiposity in WT mice.^41^

Notably, our data demonstrate that organismal response to CysR is dose-dependent. Complete cysteine depletion in our previous study induced coenzyme A depletion, unchecked thermogenesis, and rapid 30% weight loss within one week that is incompatible with life.^5^ Here we demonstrate that current sustained 70% CysR dose elicited significant weight loss that protects against age-associated weight gain. With this new and sustainable adaptive set point, metabolic functions were improved while preserving muscular function and activity. Notably, the beneficial metabolic effects were largely conserved in both sexes, despite modest sex-specific differences regarding body weight trajectories and muscle function. These findings define a window in which CysR is sufficient to induce adaptive nutrient-stress responses without compromising essential cysteine-dependent processes.

Aging is associated with increased visceral adiposity, loss of adipose plasticity, whitening of BAT, and impaired thermogenesis, which in turn impairs organismal metabolism.^42^ In addition, recent studies indicate that adipose tissue is a major organ in inflammaging.^24,43^ Our findings suggest that moderate CysR prevented these alterations by preserving BAT morphology and promoting beige remodeling of SAT, but not in VAT. This depot specificity is consistent with recent work showing that SAA restriction elicits a much stronger transcriptional response in SAT compared to VAT, suggesting a difference in depot sensitivity to amino acid restriction.^44^ Specifically, CysR shifted the adipocyte transcriptional landscape toward a youthful phenotype, with expansion of oxidative and thermogenic adipocytes, which was accompanied by reduced inflammaging and improved glucose homeostasis and endocrine adaptations.

The liver is another central node of metabolic homeostasis and a major site of age-associated dysfunction.^24^ Interestingly, we demonstrated that the liver underwent extensive adaptive remodeling, including reduced fibrosis and lipid accumulation, during aging. Despite a reduction in hepatic GSH content by approximately 35%, CysR mice maintained redox homeostasis, as evidenced by the absence of aberrant lipid peroxidation. In addition, we observed that NRF2 was not required for complete cysteine-deficiency-induced body weight loss. These data are consistent with recent murine models of hepatic GSH depletion that showed reduced hepatic lipogenesis,^45,46^ suggesting that GSH reduction induces adaptive lipid remodeling rather than overt oxidative damage, which could be of interest in the context of hepatic steatosis. Additionally, the significant induction of IGFBP2 in the liver of 70% CysR mice supports reduced IGF-1 bioavailability implicated in lifespan extension and improved hepatic homeostasis.^23^ Induction of hepatic *Igfbp2* expression is a feature shared with other pro-longevity interventions, including SAAR, caloric restriction, and deficiency in growth hormone receptor.^47–49^ This suggests cysteine restriction induces a convergence of canonical longevity-associated pathways.^50^ Moreover, in line with previous studies with protein or SAA restriction in mice and humans,^51–53^ we found that cysteine reduction, but not methionine, induced the expression of the pro-longevity hepatokine FGF21. Notably, using CTH and FGF21 double-deficient mice, we recently demonstrated that FGF21 is dispensable for cysteine-depletion-induced adipose browning.^5^

Elevated circulating cystine is consistently associated with obesity and aging in humans.^15,54^ Caloric restriction in humans lowers cysteine in adipose tissue,^5^ demonstrating that this axis is amenable to dietary modification that extends lifespan. Consistent with this, it was recently found that aging increases cystine concentration in lysosomes, which was prevented by CR and exercise in animal models.^55^ Moreover, our work complements data highlighting that intracellular cysteine excess may drive toxicity and age-associated mitochondrial dysfunction.^56^ Consistent with this, cysteine, but not methionine restriction, extended lifespan in *C. elegans*. Furthermore, the survival benefit of CysR in heat-stressed *C. elegans* was abrogated when cysteine was restored. Together, these findings underscore that cysteine availability is a previously unrecognized determinant of healthy lifespan. Notably, although cysteine depletion activates the upstream sympathetic nervous system, the downstream UCP1-independent mechanism of cysteine restriction’s adipose browning effect remains undiscovered. Expression of UCP1 in human adipose tissue is limited to discrete beige depots and is absent in white adipose depots; therefore, UCP1-independent components of the CysR-induced thermogenesis mechanism are highly relevant to future therapeutic targeting. Also, the impact of long-term lowering of the intracellular cysteine pool on lysosome-mitochondrial communication and iron-sulfur cluster function could reveal dormant pathways that impact healthspan. In terms of pro-longevity mechanisms, CysR-induced robust upregulation of LAMP2A and IGFBP2 supports the hypothesis that elevated autophagy and reduced IGF-1 bioavailability extend lifespan in animals,^38,40^ but this needs additional experiments to test causality. Lastly, although CysR but not MR promoted stress survival and lifespan of *C. elegans*, future studies in mammalian models are required to establish cysteine reduction as a mechanism of lowering disease burden and longevity.

From a translational perspective, prior efforts have focused on SAA restriction or its mimetics. As the TSP can buffer reductions in dietary methionine by sustaining endogenous cysteine synthesis, our data suggest that pharmacological inhibition of CTH together with low-cysteine, plant-based diets can potentially be tested to lower inflammaging and improve metabolic health in humans by keeping the methionine-regulated pathways intact. In summary, consistent with the association of excess cystine with biological aging and disease burden,^15^ our data provide direct evidence that selective reduction of cysteine controls immunometabolic mechanisms that promote healthy lifespan.

## RESOURCE AVAILABILITY

### Lead contact

- Requests for further information and resources should be directed to and will be fulfilled by the lead contact, Vishwa Deep Dixit.

## Materials availability

- This study did not generate new unique reagents.

## Data and code availability

- Sequencing data (liver bulk RNA-seq and adipose snRNA-seq) will be made publicly available upon publication of the manuscript.
- This paper does not report original code.
- Any additional information required to reanalyze the data reported in this paper is available from the lead contact upon request.

## ACKNOWLEDGMENTS

We thank the Yale School of Medicine’s comparative pathology research core for assistance with histological analyses and the Yale Center for Molecular and Systems Metabolism’s metabolic cage core for metabolic phenotyping. B.S. acknowledges funding from the European Research Council (ERC-2023-SyG, 101118919), the Hevolution Foundation (HF-GRO-23-1199212-35), the Deutsche Forschungsgemeinschaft (Reinhart Koselleck-Project 524088035, FOR 5504 project 496650118, FOR 5762 project 531902955, SFB 1678, SFB 1607, CECAD EXC 2030 – 390661388, DFG-ISF project 561031107, ANR-DFG project 545378328, and the DFG project grants 558166204, 540136447, 496914708, 437825591, 437407415, 418036758), the José Carreras Leukemia Foundation, DJCLS 04 R/2023, the Deutsche Krebshilfe (70114555), and the Bundesministerium für Forschung, Technologie und Raumfahrt (BMFTR) X-AGE project 02NUK105B. This study was supported, in part, by National Institutes of Health grants AG073969, P01AG051459, and U54AG079759 and Yale University Intramural funding (to V.D.D.).

## AUTHOR CONTRIBUTIONS

L.O. designed the study, performed overall experiments, prepared the figures, and wrote the first draft of the manuscript. H.H.K. performed and analyzed multi-color flow cytometry and snRNA-seq, prepared and arranged the figures, and wrote the first draft of the manuscript. P.G. and B.S. performed C. elegans experiments. A.L., Y.L., Y.H.Y., T.D., and C.G. helped with animal experiments. R.L., C.Z., M.N.A., and Y.K. assisted with snRNA-seq analysis. A.W. provided Nfe2l2 knockout mice. V.D.D. designed and supervised the study, helped interpret the data, secured funding, and wrote the manuscript.

## SUPPLEMENTAL INFORMATION

Document S1. Figures S1–S4 and Table S1.

## STAR METHODS

### Mice

All mice were on a C57BL/6J genetic background. *Cth*^-/-^ mice (C57BL/6NTac-Cth^tm1a(EUCOMM)Hmgu/Ieg^) were purchased from the European Mouse Mutant Cell Repository. All mice were housed in pathogen-free facilities in ventilated cage racks that deliver HEPA-filtered air to each cage with free access to sterile water through a Hydropac system. Mice were fed ad libitum with either CTRL diet (#511387), CysF diet (#510027), MR diet (#11474), MCR diet (#510029), 25% CysR diet (#511437), or 70% CysR diet (#511442) from Dyets. Mice were housed under a 12-h light-dark cycle with controlled humidity and temperature. All experiments were performed at Yale School of Medicine and were approved by the Institutional Animal Care and Use Committee at Yale University. Animals were randomly assigned to experimental groups, and both males and females were included in the study. *Nfe2l2*^-/-^ mice (#017009, The Jackson Laboratory) were kindly provided by Andrew Wang (Yale University) and crossed to *Cth*^-/-^mice to generate *Cth*^-/-^*Nfe2l2*^-/-^ double-knockout mice. Young (2 months) and old (22 months) C57BL/6J male mice were obtained from the National Institute on Aging (NIA) Aged Rodent Colony and housed in facilities at Yale University.

### C. elegans

All strains were maintained at 20°C as described previously unless stated otherwise.^57^ Strains used in this study were WT (N2) and *cth-1(ok3319)V*. Worms were fed either with OP50 or BW25113 *E. coli* strains. Lifespan assays were performed on nematodes *C. elegans* synchronized by alkaline hypochlorite treatment (bleach-synchronized) as described previously.^58,59^ L1 larvae were transferred to nematode growth medium (NGM) plates seeded with *E. coli* OP50, BW25113, or BW25113 derived from *cysE* or *metB* mutant strains and maintained at 20 °C for the duration of the experiment. Lifespan scoring was started from the first day of adulthood.^59^ Worms were transferred every day on a freshly seeded plate to prevent any bacterial and progeny contamination following standard *C. elegans* lifespan assay. Worms were scored as dead when they no longer responded to gentle prodding with a platinum wire, and were censored if they went missing, bagged (internal hatching), or exploded (intestinal material comes out because of cuticle rupture). For heat stress recovery survival, synchronized adult worms were exposed to an acute heat shock at 35 °C for 4 hours, followed by recovery at 20 °C for 24 hours.^22,60^ Animals were monitored daily and scored as alive or dead until all worms in the cohort had died as described above, and were transferred to freshly seeded plates every day during the fertile period to prevent confounding by progeny and maintain food quality.

### Metabolic cages

The energy expenditure, RER, activity, and food intake were monitored using Promethion Core metabolic cages (Sable Systems). Each mouse was singly housed for at least 3 days for acclimation before being placed in the individual chamber for at least 5 days of measurements. Energy expenditure was calculated based on oxygen (O_2_) consumption and carbon dioxide (CO_2_) production, which were measured every 30 minutes. An infrared beam detected mouse activity. Mice had free access to a running wheel, and food intake was measured via weight sensors placed on the food dispenser.

### Healthspan analyses

Rotarod test (Med-Associates) was performed to determine motor coordination and balance. Mice were trained for 2 consecutive days to acclimate to the movement on the machine. Each training session consisted of 3 trials of 1 minute on a slowly accelerating rod, with 30-40 minutes of rest in between. For the actual test, mice were placed on the rod, and the latency to fall was recorded for each animal. Mice were tested 3 times, and latency to fall was calculated as the average of the 3 trials.

The grip strength test is used to assess the maximal muscle strength of the forelimbs. While the mouse grasps the bar with its forepaws, its tail is gently pulled backwards. The maximum strength of the grip before grip release was recorded. Data are presented as the average measurement of three trials.

Body composition with lean and fat mass was determined by magnetic resonance imaging (EchoMRI, Echo Medical Systems). For the analysis, each mouse was placed into an acrylic tube with breathing holes and inserted into the machine for approximately 90 seconds.

For the glucose tolerance test, mice were fasted overnight before being injected intraperitoneally with glucose solution (2g/kg of body weight). For the insulin tolerance test, mice were fasted for 5 hours before being injected intraperitoneally with insulin (0.5U/kg of body weight). Tail vein blood was collected to measure glycemia with a glucometer.

### Tissue collection

Immediately after dissection, tissues were either digested with collagenase, snap-frozen for further analysis, or drop-fixed in 10% formalin for 24 hours for histological analysis. Blood was collected by cardiac puncture and allowed to clot for 2 hours at room temperature. Serum was collected after centrifugation for 20 minutes at 3,000 rpm.

### Serum and tissue parameter measurements

Serum FGF21 and GDF15 levels were determined by ELISA (R&D). Leptin, MCP1, TNFα, and IL-6 were quantified by a 6-plex magnetic bead panel for Luminex (Mouse metabolic hormone panel, Sigma). For CoA and total GSH quantification, frozen biopsies were homogenized in ice-cold PBS supplemented with EDTA and protease and phosphatase inhibitors. Protein concentration was determined by colorimetric assay (Bio-Rad) before filtering samples on 10 kDa Amicon ultracentrifugal filters (Millipore). CoA (Abcam) and total GSH (Cayman) levels were determined by colorimetric assays, following manufacturers’ recommendations.

### Histological analysis

Tissues were embedded in paraffin and sectioned into 5-10 μm thick sections. Sections were stained with hematoxylin and eosin according to standard procedures. For UCP1 staining, sections were probed with anti-UCP1 antibody (1:100, Abcam), followed by goat anti-rabbit HRP-conjugated antibody (DAKO). Slides were developed with DAB peroxidase (Vector Laboratories) and counterstained with hematoxylin. Sections were imaged with a Keyence microscope. Adipocyte size was quantified with Adiposoft macro on ImageJ (Fiji).^61^

### Western blot

Proteins were extracted from frozen tissues in ice-cold RIPA buffer (Sigma) supplemented with 1x protease and phosphatase inhibitors (Sigma). Protein concentration was determined with a DC Protein Assay (Bio-Rad). Proteins were separated on NuPAGE 4-12% Bis-Tris polyacrylamide gels (Invitrogen) and then transferred onto nitrocellulose membranes with a semi-dry transfer system (Bio-Rad). Membranes were blocked in 5% non-fat dry milk in TBST (Bio-Rad) and incubated with primary antibody overnight. HRP-conjugated anti-mouse (Invitrogen) or anti-rabbit (Invitrogen) antibodies were used accordingly. Membranes were visualized with Pierce ECL or SuperSignal West Pico Chemiluminescent substrate, with a ChemiDoc MP Imaging System (Bio-Rad). Images were quantified with ImageJ software (FIJI). The following antibodies were used: CTH (Novus, H00001491-M03), UCP1 (Abcam, ab10983), 4-HNE (Thermo, MA5-27570), IGFBP2 (Proteintech, 11065-3-AP), LAMP2A (Abcam, ab125068), HSP40 (Cell Signaling, CS4868), HSP90 (Cell Signaling, CS4874) and ACTIN (Cell Signaling, CS4967).

### Flow cytometry

The spleen was harvested, and its weight was recorded. The spleen was put on a 70-μm cell strainer and minced using a syringe plunger. Isolation medium (RPMI1640 with 10% fetal bovine serum) was added to a cell strainer, and samples were centrifuged at 650 xg for 5 min at 4 °C. The pellet was resuspended in isolation media and filtered through a 40-μm cell strainer. Cells were then centrifuged again, and the pellet was used for further analysis. Isolated splenocytes were incubated with anti-mouse CD16/CD32 (BD Biosciences) Fc-blocker for 5 min on ice, followed by staining with various antibodies: BV711-CD45 (30-F11) (Biolegend), FITC-CD4 (RM4-5) (eBioscience), APC-CD8a (53-6.7) (Invitrogen), PE-CD44 (IM7) (eBioscience), BV605-CD62L (MEL-14) (Biolegend), FITC-F4/80 (BM8) (Biolegend), APC-CD11c (N418) (Biolegend), APC780-CD206 (MR6F3) (Invitrogen), or PerCP-Cy5.5 CD11b (M1/70) (Invitrogen) for 40 min at 4 °C in the dark. Cells were then washed with ice-cold saline twice, and to identify dead cells, the LIVE/DEAD^™^ fixable aqua dead cell stain kit for 405 nm excitation (Thermo) was added for 20 min on ice in the dark. After another wash with saline, cells were fixed with 1% paraformaldehyde and read with an LSRII flow cytometer (BD). All the data were processed by FlowJo software (version 10; FlowJo LLC).

### Quantitative RT-PCR

Total RNA was isolated from frozen tissues with RLT buffer supplemented with beta-mercaptoethanol, followed by the RNeasy kit (Qiagen) according to the manufacturer’s instructions. Complementary DNA was synthesized with the iScript cDNA synthesis kit (Bio-Rad). Quantitative PCR reactions were performed with Power SYBR Green detection reagent (Thermo Fisher) and sequence-specific primers (Table S1) on a QuantStudio 6 Pro Real-Time PCR System (Thermo). *18s* was used for normalization of relative expression levels.

### Liver RNA-sequencing

RNA was extracted from snap-frozen liver samples as previously detailed. Raw sequencing reads were processed using the nf-core/rnaseq pipeline (v3.18.0)^62^ with default parameters and the STAR-Salmon quantification route. Reads were aligned to the mouse reference genome (GRCm38/mm10) using STAR^63^, and transcript-level quantification was performed with Salmon^64^. The pipeline was executed with Nextflow (v24.10.3)^65^ using Singularity^66^ containers. Transcript-level abundance estimates were imported and summarized to the gene level using tximport (v1.32.0)^67^ with transcript-to-gene mappings from Ensembl release 98 (*M. musculus* GRCm38, AnnotationHub ID: AH89211). A DESeqDataSet was constructed, and differential expression analysis between 70% CysR (*n* = 4) and CTRL groups (*n* = 3) was performed using DESeq2 (v1.44.0)^68^. Genes with fewer than 10 counts in at least 3 samples were excluded before analysis. Log2 fold-change estimates were shrunk using the apeglm^69^ method. Genes with a Benjamini–Hochberg adjusted p-value < 0.05 and an absolute log2FC > 2 were considered differentially expressed. Results were visualized with volcano plots and heatmaps of variance-stabilizing transformed expression values.

For pathway analysis, fgsea^70^ was performed, with a minimum of 15 and a maximum of 500 genes in a pathway, and with 1 million permutations. We retrieved GO terms for *M. musculu*s from category M5 (subcategory GO:BP, GO:CC and GO:MF), canonical pathways from category 2, subcategory CP, and hallmark gene sets for *M. musculus* from category H (MSigDB v2024.1). Redundant significant terms were eliminated. When two terms had more than 60% of genes in common in their leading gene sets, the term with the lowest normalized enrichment score was considered redundant and eliminated. For gene signatures, the following gene sets were used as reference: Aging hepatocyte gene signature^31^, SenMayo^32^, and FerrDb^37^.

### SnRNA-seq of SAT and BAT

Snap-frozen tissues were weighed and pooled (*n* = 3/group), and nuclei were isolated following the previous literature with minor modifications.^71,72^ Briefly, approximately 200 mg of frozen tissues were minced with scissors on ice in a 20 mL scintillation vial containing 500 µl of isolation buffer (0.5 M sucrose, 1 M HEPES, 150 mM MgCl2, 2 M KCl, 1 % Triton X-100, 0.1 M DTT, and 40,000 U/ml murine RNase inhibitor in water). Minced tissues were then moved to a pre-cooled 1.5 mL DNA LoBind tube and homogenized with a hand motor with a pestle for 10 sec on ice. After that, tissues were filtered with a 100 µm strainer, moved to a 5 ml DNA LoBind tube, and centrifuged at 1,000 xg for 10 min at 4 °C. The supernatant was carefully discarded, and the pellet was resuspended and transferred to a new pre-cooled 5 mL DNA LoBind tube. Isolation buffer was added to 3 mL, and tubes were centrifuged at 500 xg for 10 min at 4 °C. After carefully discarding the supernatant, the pellet was resuspended in 100 ul of resuspension buffer (5 % bovine serum albumin and 40,000 U/ml murine RNase inhibitor in saline). Nuclei were then filtered with a pre-wet 40 µm Flowmi tip strainer and were counted with trypan blue. The concentration of isolated nuclei was between 1,200 and 1,900 nuclei per microliter. Nuclei processing was conducted with the Chromium Next GEM Single Cell 3’ Reagent Kits (v4). The targeted number of nuclei was 20,000 per sample. Sample barcoding and library preparation were conducted according to the manufacturer’s instructions (10x Genomics). HiSeq3000 (Illumina) was used to sequence the libraries at 50,000 read pairs per nucleus depth.

Single-cell transcriptomes from mouse BAT and SAT samples under 70% CysR and CTRL conditions were sequenced using the 10X Genomics Chromium platform. Raw sequencing data were processed using the standard Cell Ranger pipeline. Count matrices were then loaded and merged into a single Seurat object.^73^ For quality control, cells were filtered based on tissue-specific thresholds: cells with > 2,500 detected genes (BAT) or > 5,000 detected genes (SAT), mitochondrial content ≥ 3%, or ribosomal content ≥ 5%, and miQC.keep^74^ label of “discard” were removed. Doublets were predicted and removed using DoubletFinder^75^ with parameters PCs = 1:20, pN = 0.25, and optimal pK, assuming the doublet formation rate is 5%. Genes detected in fewer than 5 cells across all samples of a given condition were excluded. The samples are then merged into one dataset and processed using log-normalization. Variable features were identified using the mean.var.plot method with a mean cutoff of 0.0125 to 5 and a dispersion cutoff of > 0.5. Reads are scaled by regressing out mitochondrial RNA percentage and number of features. All variable features were used for downstream analysis. Principal component analysis is performed using default parameters, and batch effects are corrected using Harmony^76^ by regressing over the samples. The top Harmony-corrected principal components were used for non-linear dimension reduction and clustering. Then, the dataset was divided into the adipocyte and non-adipocyte subsets for annotation and downstream analysis, where clustering resolutions of 0.2 and 0.1 were used for the SAT and BAT subsets, respectively. For each BAT and SAT cluster, differential expression analysis across conditions was performed using the FindMarkers function with default parameters. Genes with adjusted p-value < 0.01 and log2 fold change greater than 0.5 were selected for GO enrichment analysis using clusterProfiler^77^. Gene expression module scores were generated with the AddModuleScore function of the Seurat package. Gene sets were retrieved from GO:BP data sets. Only terms containing between 15 and 200 genes, and related to oxidative phosphorylation, thermogenesis, fatty acid oxidation, and autophagy, were kept for subsequent analysis. Gene sets were intersected with genes expressed in at least 10 nuclei. Module scores were computed on all adipocytes and at the cluster-level by computing mean scores across nuclei. Statistical significance of differences between conditions was assessed using two-sided Wilcoxon rank-sum tests.

### Statistical analysis

Investigators were not blinded during data collection. Data were analysed in R (v4.3.0) and GraphPad (v10). Statistical differences between groups were calculated using unpaired two-tailed t-tests when comparing two groups or two-way analysis of variance with Sidak’s correction for multiple comparisons. Data distribution was assumed to be normal, but this was not formally tested. For the survival analysis of *C. elegans*, the log-rank (Mantel-Cox) method was used to test the null hypothesis (two-sided hypothesis test) in Kaplan-Meier survival analysis. P-value < 0.05 was considered significant. Sample sizes were not statistically pre-determined and are similar to those previously reported in the field. Mice with abnormal health or reaching a humane endpoint were removed from the study.

---

**Figure S1.**
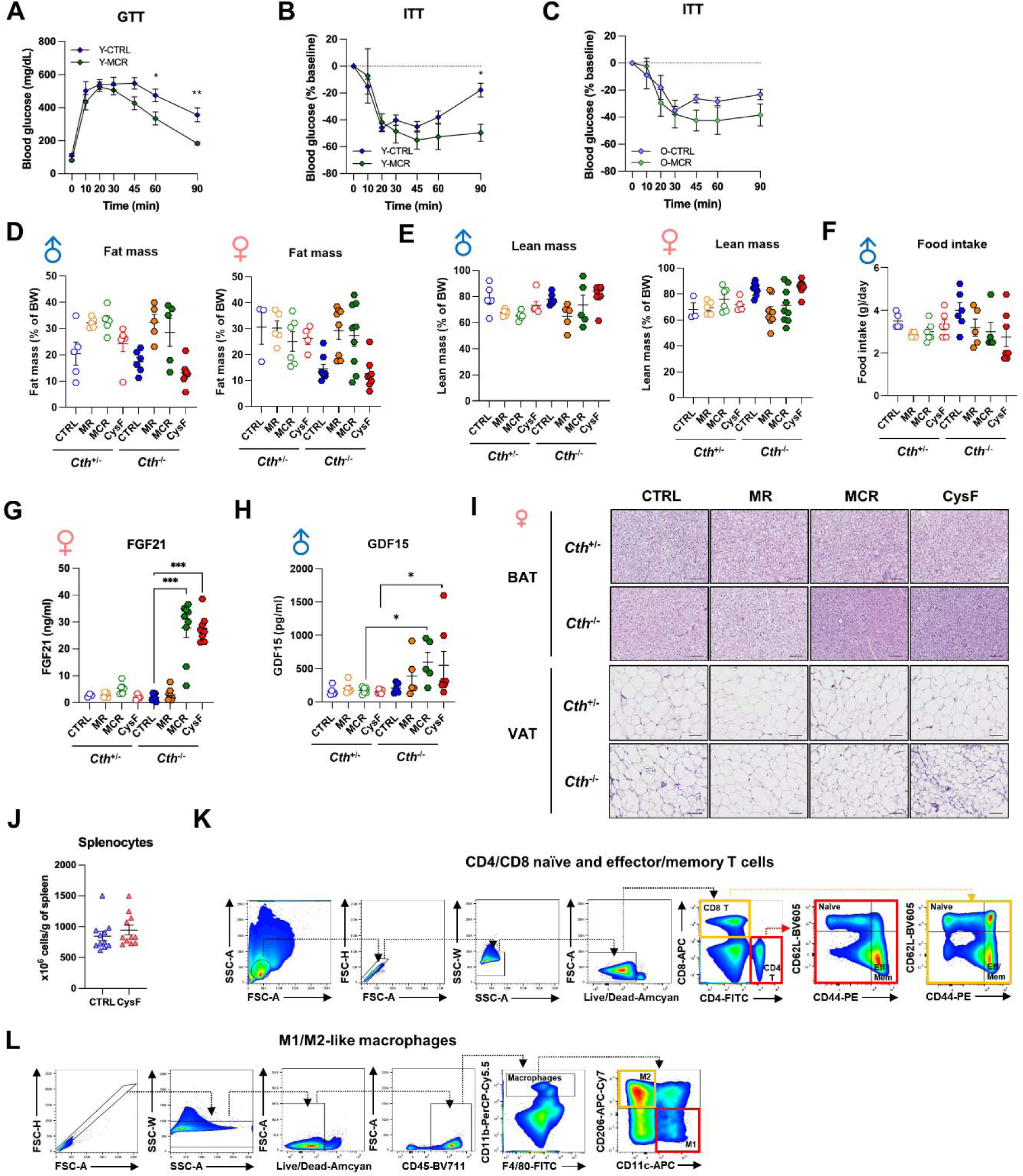
Cysteine depletion, but not methionine restriction, drives mechanisms that induce adipose browning. Related to Figure 1. (A-C) Young or old male mice were fed either the control diet (CTRL) or the methionine-and cysteine-restricted diet (MCR) for 12 weeks (*n* = 4-5/group). (A) Glucose tolerance test in young mice. (B and C) Insulin tolerance tests in young (B) and old (C) mice. (D-I) Adult *Cth*^+/-^ and *Cth*^-/-^ mice were fed a CTRL, MR, MCR, or CysF diet for 4 days (*n* = 5-7/group for male and 3-9 for female mice). (D) Fat mass. (E) Lean mass. (F) Food intake in male mice. (G) Serum FGF21 levels in female mice. (H) Serum GDF15 levels in male mice. (I) Representative H&E staining of BAT and VAT of female mice. Scale bars = 50 µm. (J) Total splenocyte count (*n* = 11 for CTRL, 12 for CysF). (K and L) Representative gating strategy for flow cytometry analyses of splenic naïve and effector/memory T cells (K) and M1-or M2-like macrophages (L). Data are presented as mean±SEM. Unpaired two-tailed t-tests (A-C and J) or Two-way ANOVA with Sidak’s correction for multiple comparisons (D-H) were performed. *P <0.05, **P <0.01, and ***P <0.001.

**Figure S2.**
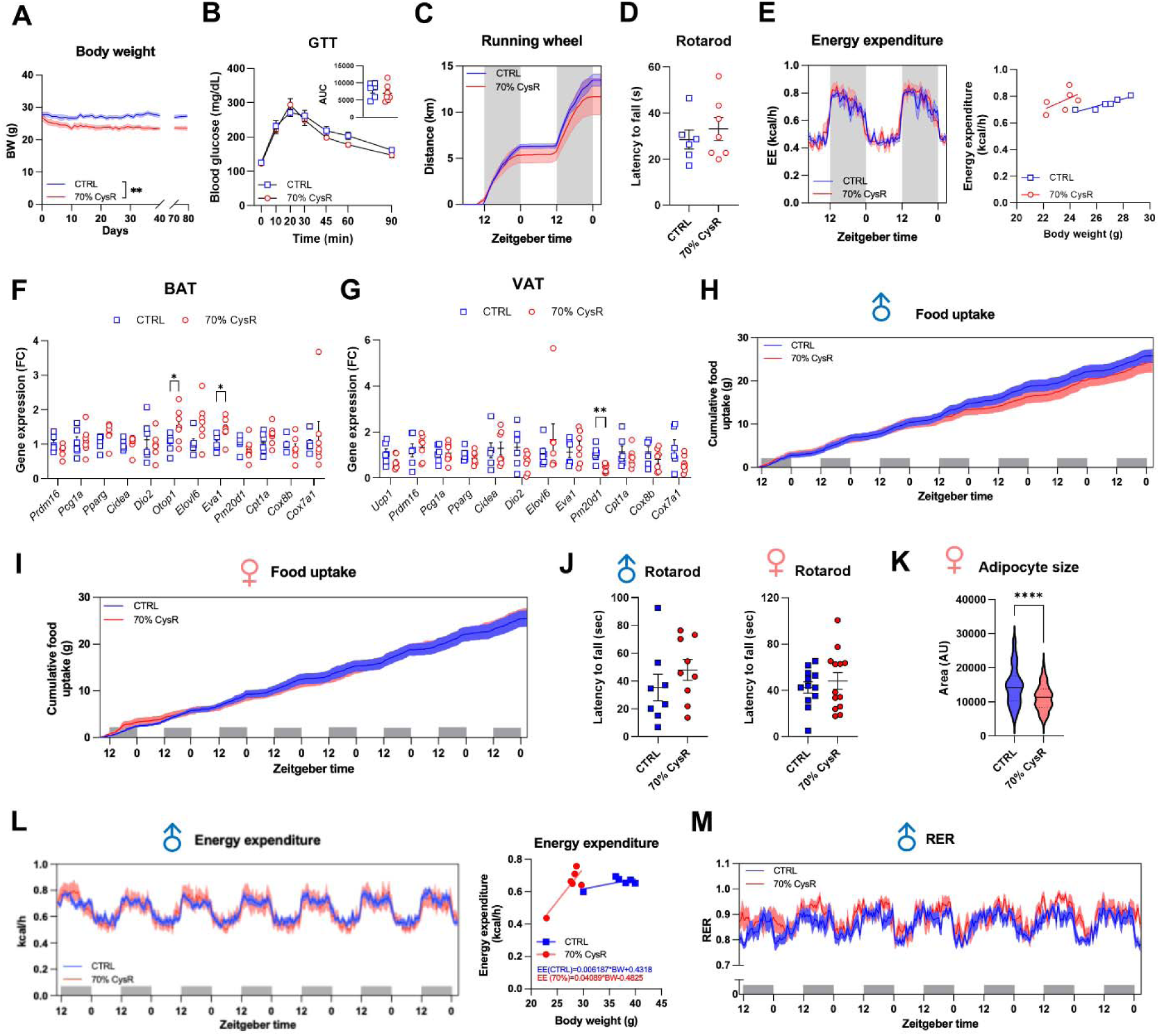
Healthspan analysis of 70% CysR young and aged CTH KO mice. Related to Figure 3. (A-H) 3-month-old *Cth*^-/-^ mice were fed a CTRL or 70% CysR diet for 11 weeks (*n* = 6-7/group). (A) Body weight changes. (B) GTT. (C) Cumulative distance on the running wheel (*n* = 6/group). (D) Rotarod test. (E) Energy expenditure (*n* = 6/group). (F and G) qRT-PCR analyses of BAT (F) and VAT (G). (H-M) 8-to 10-month-old *Cth*^-/-^ mice were fed a CTRL or 70% cysteine-restricted diet for 10 months (*n* = 13-25/group). (H and I) Cumulative food intake for 1 week in male (H) and female (I) mice (*n* = 6/group). (J) Rotarod test in male (left) and female (right) mice (*n* = 8-12/group). (K) SAT adipocyte size calculation in female mice. (L) Energy expenditure in male mice (*n* = 6/group). (M) RER in male mice (*n* = 6/group). Data are presented as mean±SEM. Unpaired two-tailed t-tests were performed. *P <0.05, **P <0.01, and ****P <0.0001.

**Figure S3.**
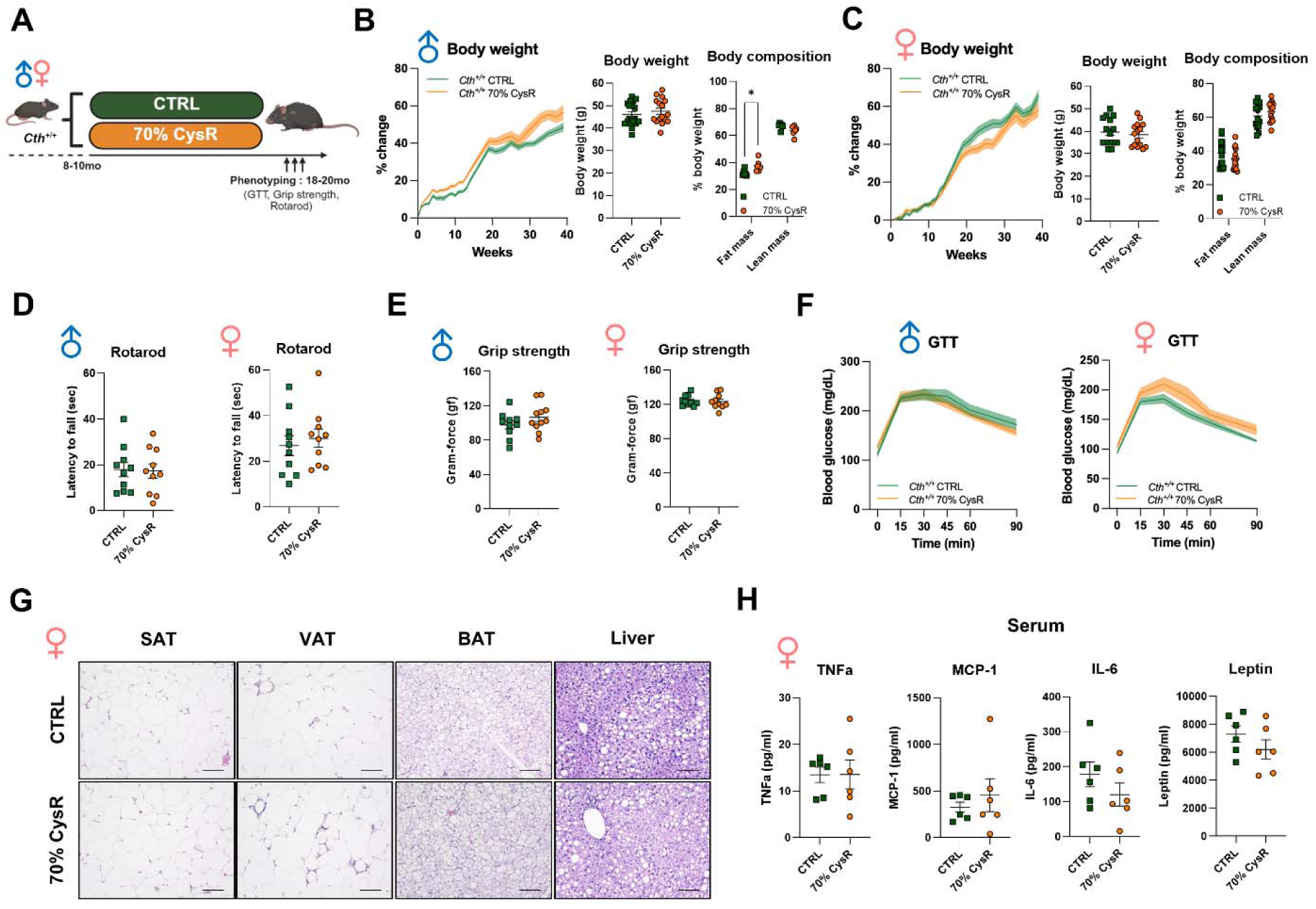
Dietary 70% cysteine restriction does not impact healthspan in WT mice. Related to Figure 3. (A-H) 8-to 10-month-old *Cth*^+/+^ mice were fed a CTRL or 70% cysteine-restricted diet for 10 months (*n* = 25-37/group). (A) Schematic outline. (B and C) Body weight changes and body composition in male (B) and female (C) mice. (D) Rotarod test in male (left) and female (right) mice (*n* = 10/group). (E) Grip strength test in male (left) and female (right) mice. (F) GTT in male (left) and female (right) mice (*n* = 10-11/group). (G) Representative H&E staining of SAT, VAT, BAT, and liver tissues of female mice. Scale bars = 50 µm. (H) Serum levels of TNF-α, MCP-1, IL-6, and leptin in female mice (*n* = 6/group). Data are presented as mean±SEM. Unpaired two-tailed t-tests were performed. Figure S3A was created with Biorender.com. *P <0.05.

**Figure S4.**
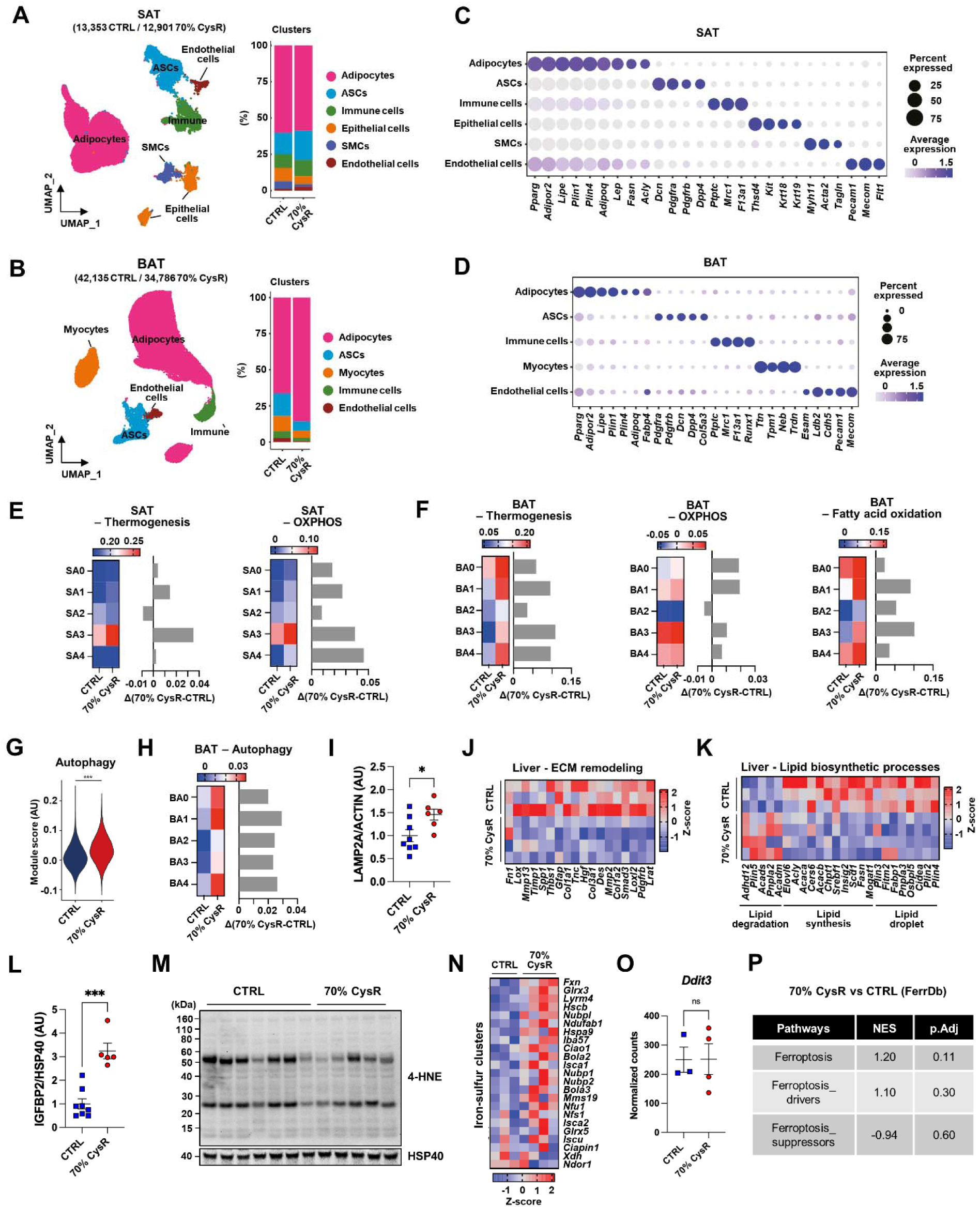
Chronic cysteine restriction induces metabolic reprogramming that favors healthspan. Related to Figure 4. (A-F) SnRNA-seq of SAT and BAT from 8-to 10-month-old *Cth*^-/-^ mice fed a CTRL or 70% CysR diet for 10 months. (A and B) UMAP for all clusters in SAT (A) and BAT (B). (C and D) Marker genes for each cluster in SAT (C) and BAT (D). (E and F) Scoring analysis for indicated pathways in each adipocyte subcluster in SAT (E) and BAT (F). (G and H) Scoring analysis for the autophagy pathway of the whole adipocyte fraction (G) and each adipocyte cluster (H) of BAT. (I) Densitometry analysis of LAMP2A western blot of BAT (*n* = 6-8/group). (J and K) Heatmaps for the genes involved in extracellular matrix remodeling (J) and lipid biosynthetic process (K) from the liver RNA-seq (*n* = 3-4/group). (L) Densitometry analysis of IGFBP2 western blot of liver (*n* = 5-8/group). (M) Representative 4-HNE western blot of the liver (5-7 blots/group). (N) Heatmap for the genes involved in the iron-sulfur cluster pathway from the liver RNA-seq (*n* = 3-4/group). (O) Normalized counts of *Ddit3* from liver RNA-seq (*n* = 3-4/group). (P) Normalized enrichment scores of ferroptosis-related gene sets queried from FerrDb in liver RNA-seq were presented with adjusted p-values. Data are presented as mean±SEM. Unpaired two-tailed t-tests were performed. *P <0.05 and ****P <0.0001. ns, not significant.

**Table S1.**
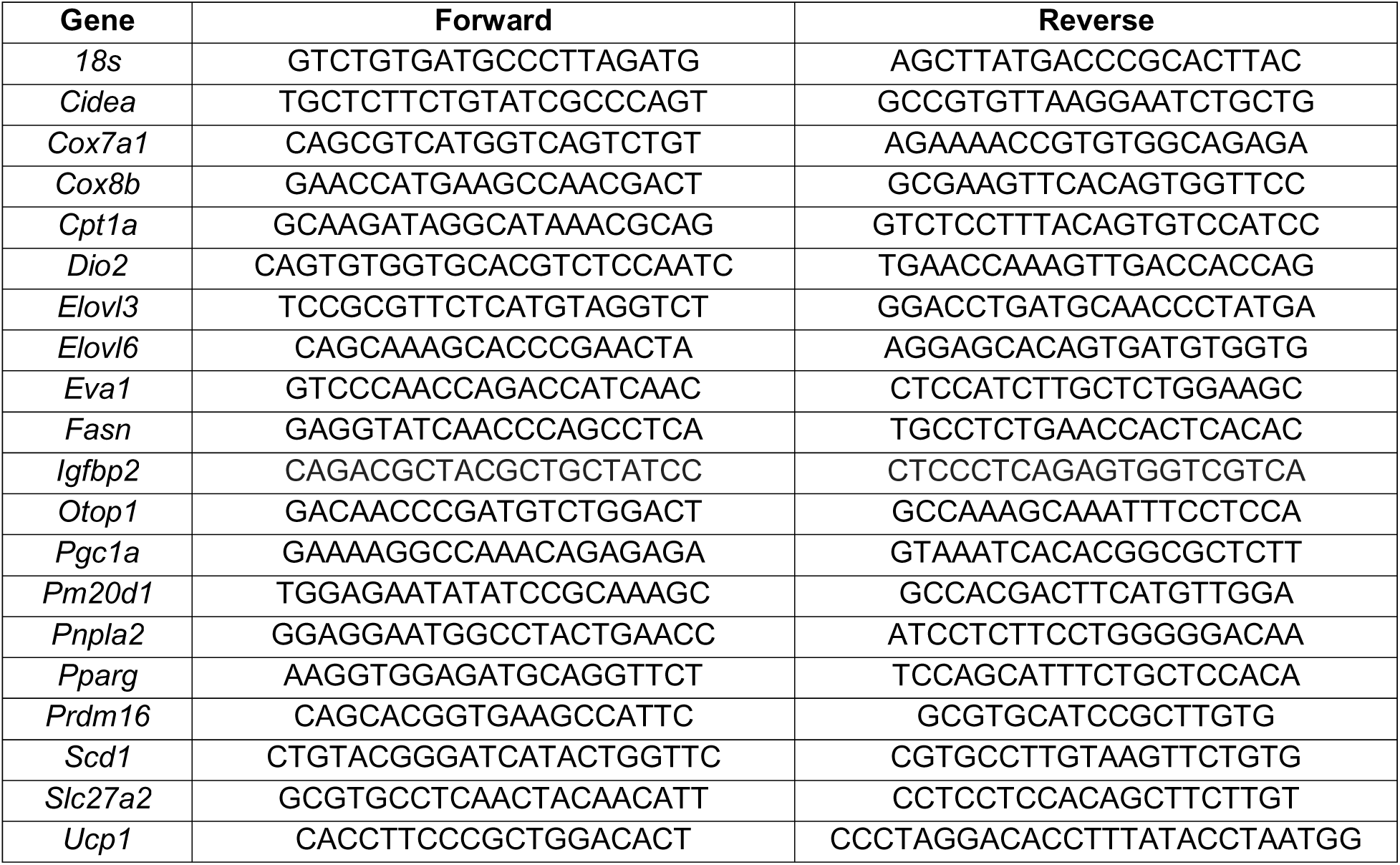
qRT-PCR primers. Related to STAR Methods.

